# Topological Feature Extraction and Visualization of Whole Slide Images using Graph Neural Networks

**DOI:** 10.1101/2020.08.01.231639

**Authors:** Joshua Levy, Christian Haudenschild, Clark Barwick, Brock Christensen, Louis Vaickus

## Abstract

Whole-slide images (WSI) are digitized representations of thin sections of stained tissue from various patient sources (biopsy, resection, exfoliation, fluid) and often exceed 100,000 pixels in any given spatial dimension. Deep learning approaches to digital pathology typically extract information from sub-images (patches) and treat the sub-images as independent entities, ignoring contributing information from vital large-scale architectural relationships. Modeling approaches that can capture higher-order dependencies between neighborhoods of tissue patches have demonstrated the potential to improve predictive accuracy while capturing the most essential slide-level information for prognosis, diagnosis and integration with other omics modalities. Here, we review two promising methods for capturing macro and micro architecture of histology images, Graph Neural Networks, which contextualize patch level information from their neighbors through message passing, and Topological Data Analysis, which distills contextual information into its essential components. We introduce a modeling framework, *WSI-GTFE* that integrates these two approaches in order to identify and quantify key pathogenic information pathways. To demonstrate a simple use case, we utilize these topological methods to develop a tumor invasion score to stage colon cancer.

## 1. Introduction

Large-scale architectural motifs and repetitive patterns of functional tissue sub-units (eg. cells, connective tissue, extracellular matrix) form the basis of histopathology. While normal tissue is relatively homogenous, cancer contains disordered structures / phenotypes that reflect driving genetic alterations. As neoplastic transformation progresses, the extent of infiltration and destruction of normal tissue is used to grade and stage cancers. Practitioners of histopathology are thus highly sensitive to disruptions in normal structure. A wide variety of computational methods have been developed to augment traditional histological inspection^1^ by reducing time and personnel costs associated with manual slide screening. These emerging techniques have also demonstrated the potential for identifying novel disease pathways and previously unrecognized morphologies.

Deep learning has been particularly successful in digital pathology ^2^. In comparison to prior modeling techniques that use handcrafted features, deep learning applies parameterized filters and pooling mechanisms via convolutional neural networks (CNN) to capture and integrate lower level image features into successively higher levels of complexity ^3^. These approaches have been used to automatically stage liver fibrosis ^4^, identify morphological features correspondent with somatic alterations^5^, assess urine slides for bladder cancer^6^, and circumvent costly chemical staining procedures^7,8^, amongst many others^9^. Many research groups are developing high-throughput clinical pipelines to take advantage of these healthcare technologies. Validating and scaling these technologies is essential for successful deployment^10^.

As a result of the gigapixel resolution of Whole Slide Images (WSI), which contain a diverse range of tissue and morphological features, researchers typically must partition the WSI into smaller sub-images. These sub-images are then evaluated separately via the deep learning model for classification or segmentation tasks, from which their results may be aggregated for slide-level inferences. Aggregation via a CNN incorporates excessive whitespace and places unnecessary dependence on the orientation and positioning of the tissue section^11^. Alternatively, a ‘bag of images’ approach can be taken, in which patch representations are aggregated using autoregressive or attention-based mechanisms to generate a whole slide representation, ignoring non-tissue regions^12–14^. These integrative approaches may be highly stochastic and insufficiently reproducible / reliable to be properly integrated into the clinic or with other omics-based modalities. These methods may additionally undervalue the higher order context between a patch and its immediate neighbors which may be vitally important to the targeted prediction.

Graphs are mathematical constructs that model pairwise relationships between entities. Accordingly, graphs are well suited to model dyadic relationships between single patches (nodes) in a WSI as defined by their spatial distance/correlation (edges). Graph Neural Networks (GNN) have been developed to encapsulate information from adjacent tissue regions/cells in order to inform the representation of the current patch of interest. GNN naturally capture the intermingling of various tissue sub-compartments while remaining permutationally invariant (the ordering/rotation of patches on slides does not impact prediction). While square-grid convolutions over WSI sub-images propagate information within a fixed neighborhood of patches and require consistent ordering of patches ^11^, GNN relax the convolutional operator to aggregate information across an unfixed number of neighbors to update the patch-level embedding^15^.

Prior GNN research on WSIs center graph nodes on cells under the assumption that cell-cell interactions are the most salient points of information^16^. However, this approach underappreciates the diagnostic/prognostic information conveyed by tissue macro-architectural structures. Constructing cell-centered graphs are limited by cell detection accuracy (a surprisingly difficult problem) and more importantly, incorporating all cells in a graph model is subject to complexity constraints. Despite these potential limitations, there remain numerous techniques to study WSI using GNN at various scales ^17^. Here, we seek methods to explain graph convolution results post-hoc to elucidate mechanisms by which tissue regions interact.

Topological Data Analysis (TDA) quantifies the underlying shape and structure of data by collapsing persistent topological structures ^18^. TDA is well-suited for summarizing Whole Slide Graphs (WSI fitted by a GNN; e.g. WSG) to identify and relate key tissue architectures, regions of interest, and their intermingling. However, the sheer quantity, complexity, and dimensionality of histology data makes interpretation challenging. A recently developed TDA-tool, Mapper^18^, alleviates this issue by providing a succinct summary of high-dimensional data to elucidate obscured relationships. Mapper projects the data to a lower dimensionality, packs the data into overlapping sets, which are then clustered to form a simplified, easily interpretable graph. Unlike pooling approaches that are built into the deep learning model and must be pre-specified, Mapper is generalizable and can be configured to study WSI information at multiple resolutions after fitting a GNN model. These models have the capacity to provide higher order descriptors of information flow for any GNN model, greatly simplifying analysis. Abstractions can then be analyzed to learn new disease biology through interrogation of patch-level embeddings. While TDA methods have previously been applied to high dimensional omics data^19–21^ and histopathology images^22,23^, to our knowledge, there have been no applications of TDA methods to GNN models fit on histological data, where these methods may be of great benefit.

Colorectal Cancer (CRC) is a common cancer with approximately 150,000 new cases annually in the United States and an estimated 63% 5-year survival rate. CRC most commonly arises from dysplastic adenomatous polyps with somatic alterations in the APC pathway or the mismatch repair (MMR) pathway^24^. The Colon is divided into distinct layers including epithelium, lamina propria, submucosa, muscularis propria, pericolic fat, and serosa (in certain anatomical locations). Tumor staging comprises tissue and nodal stages with higher numerals indicating a greater depth of invasion and greater number of lymph nodes (LN) involved by the tumor, respectively.

We present a Whole Slide Image GNN Topological Feature Extraction workflow *(WSI-GTFE)*^*^, for applying topological methods to interrogate a WSI GNN fit and demonstrate its utility in determining colon cancer stage. As a simple use case for these methods, we apply Mapper to quantitate the degree of tumor invasion into deeper sub-compartments of the colon and corroborate these tumor invasion scores (TIS) with disease staging to form an interpretable predictive score. The demonstration featured in this paper outlines some of the many potential applications of topological methods in the analysis of WSI GNN models.

## 2. Materials and Methods

### 2.1. Data Acquisition and Processing

We selected Colon (n=172) and accompanying Lymph Node (n=84) resection slides from 36 patients at Dartmouth Hitchcock Medical Center. Briefly, samples were grossed, embedded in paraffin blocks, sliced into five-micron sections and scanned using the Leica Aperio-AT2 scanner at 20x, stored in SVS image format, from which our in-house pipeline *PathFlowAI*^10^ was utilized to extract and preprocess the slides into NPY format. A board-certified pathologist provided coarse segmentation maps in the following categories: 1) epithelium, 2) submucosa, 3) muscularis propria, 4) fat, 5) serosa, 6) debris, 7) inflammation, 8) lymph node, and 9) cancer. We extracted 2.69 million 256×256 pixel patches correspondent to these slides.

### 2.2. Overview of Framework

The *WSI-GTFE* framework (**Figure 1**), provides methods to summarize the intermingling of tissue sub-compartments via a two-stage CNN-GNN model, followed by utilization of TDA methods:

1. Learning patch-level CNN embeddings and constructing spatial adjacency graph (**Figure 1A**)
2. Contextualizing patch level embeddings via an unsupervised or supervised GNN (**Figure 1B**)
3. Optionally refining the patch level embeddings through estimation of uncertainty in patch-level classification tasks (**Figure 1C**)
4. Applying Mapper to pool patches into overlapping Regions of Interest (ROI) (**Figure 1D-E**)
5. Estimating the degree of information flow and intermingling between the regions (**Figure 1F**)
6. Optionally using measures of information flow as additional markers for clinical or molecular associations (**Figure 1G**)

**Fig. 1.**
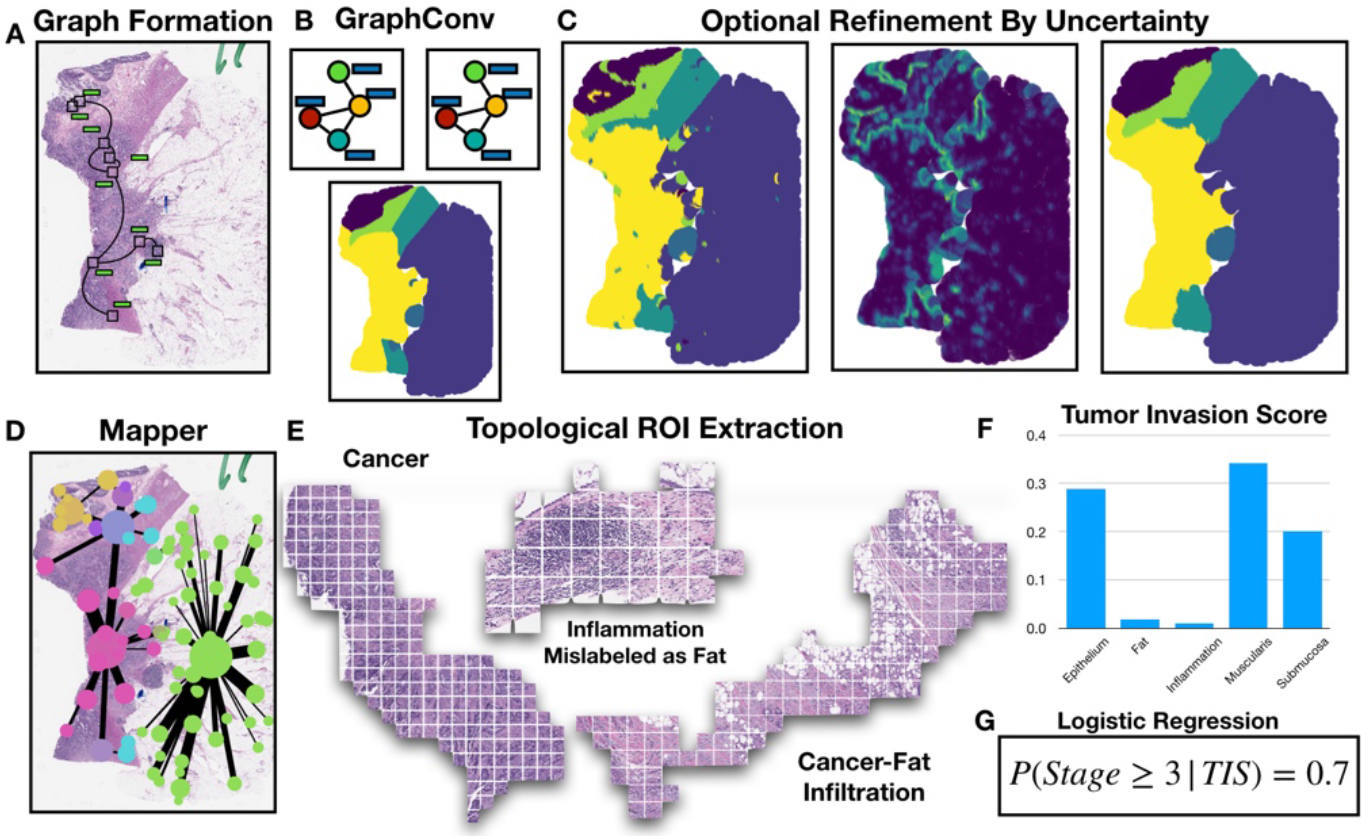
WSI-GTFE Framework: a) patch-level CNN embeddings extracted using *PathFlowAI* form graph via their spatial adjacency; b) targets (eg. colon sub-compartments) predicted using successive applications of graph convolutions; c) highly uncertain regions (middle) from noisy prediction map (left) may be reassigned (right); d) Mapper summarizes GNN embeddings over WSI as a graph; e) Meaningful histology (ROI) captured as Mapper graph nodes; f) Functional relationships between Cancer and other ROI, weighted edges Mapper graph, mined to form *TIS* vector; g) *TIS* used in prediction model to form interpretable staging score (odds ratios and log-odds probability), demonstrates type of relationships that may be extracted using TDA

#### 2.2.1. Estimation of Patch-Level Embeddings

A WSI (an RGB array on the order of 100,000 pixels in any spatial dimension) 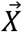, is comprised of a collection of sub-images, 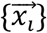. A neural network maps each sub-image to a low dimensional embedding or representation, 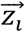, via the following mapping 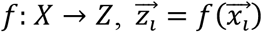. Patch-level features may be extracted using pretrained CNNs such as ImageNet, which has learned a huge collection of convolutional filters and features correspondent to 1000 common objects such as dogs, cats and birds ^25^. Features may also be acquired using unsupervised approaches such as variational autoencoders (VAE)^26^ or self-supervised techniques such as contrastive predictive coding (CPC)^14^ or SimCLR^27^. Finally, patch-level features may be learned after pretraining on histology targets of interest, such as classified objects or ROI. We utilized both an ImageNet-pretrained CNN as well as a CNN we pretrained for tissue sub-compartment classification task, generating two separate sets of patch-level embeddings for comparison.

#### 2.2.2. Contextualizing Patch-Level Embeddings via GNN

Graphs are represented via following expression *G* = *(V, E, A, X)*. The set of nodes/patches or vertices *V* are related to each other via edgelist *E*. Alternatively, the edgelist may be represented by a sparse adjacency matrix *A*, of which binary indicators *A*_*ij*_ depict a relationship between node *i* and node *j*. Node/patch-level embeddings or features are represented by attribute matrix *X*. A WSI may be encoded as a graph by storing patch level embeddings (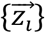, index *i* for select patch) in the attribute matrix *X* (m patches by n embedding dimensions) and recording spatial adjacency (via a k or radius nearest neighbors) of all patch coordinates as *A*. GNN utilize message passing operations to update the embeddings of nodes by their neighbors via the following convolution operation ^28^:

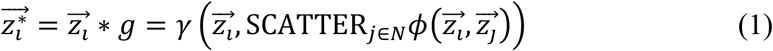

The embeddings of the neighbors of patch *i*, in neighborhood *N*, are themselves updated via some parameterized functional ϕ, which is scattered in parallel across GPUs, and then aggregated to update the embedding of patch *i* via the parameterized operation γ. Information from neighboring patches are passed as such. Multiple applications of these convolutions expand the neighborhood from which information is propagated. Additional pooling mechanisms, *AGG*, such as DiffPool or MinCutPool^29,30^, serve to aggregate the patch level representations into cluster or slide-level representations:

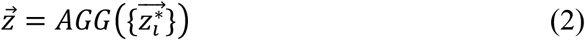

There exist multiple modeling objectives for updating these embeddings, which include: 1) node-level classification, where 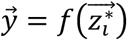, trained via the cross-entropy loss, 2) unsupervised node-level measures such as Deep Graph Infomax^31^ and spectral clustering objectives, and 3) graph-level supervised, eg. 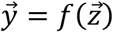, or 4) graph-level self-supervised objectives. For demonstration purposes, we learn patch-level classification of colon sub-compartments and predict these sub-compartments on held-out slides after initialization of an adjacency matrix of patches, which could be used to pretrain whole-slide level objectives. From the fitted GNN model, intermediate patch-level or cluster-level (when applying pooling operations) embeddings may be extracted for further analysis. While we constructed WSG from the spatial adjacency of patches in this work, this *WSI-GTFE* method is agnostic of WSG creation approach. These graphs also may be built using cell / nucleus detection methods, though such methods are beyond the scope of this work.

#### 2.2.3. Optional Refinement of Patch-Level Predictions

Graph convolutions aim to contextualize patches with their neighbors and as such are able to smooth the map of predictions across a slide. However, small deformities in an otherwise homogenous decision map, (for instance, pockets of inflammation that were not captured by the pathologist’s relatively coarse annotations), may be a source of signal noise. To further smooth the classification map of patches across a slide, assigned patch-level labels may be refined using label propagation techniques. Dropout^32^ methods randomly set predictors at a particular neural network layer to 0 with a certain probability, while DropEdge^33^ randomly prunes edges of a graph, which in this case corresponds to the adjacency matrix of the WSI. While both of these techniques have been utilized to improve the generalization of graph neural networks through perturbations to the input data and intermediate outputs, applications of these techniques during prediction may be used to make multiple posterior draws of a patch-level categorial distribution for class label assignment. Both the variance of the predictive posterior distribution after numerous posterior draws and the entropy in the class labels after averaging the results for a sample after application of SoftMax layer may be used as estimates of uncertainty in prediction^34^. Nodes that exhibit high uncertainty may be pruned and the remaining class labels may be propagated to the unlabeled patches.

#### 2.2.4. Application of Mapper to Extract Regions of Interest

Once a GNN model has been fit, post-hoc model explanation techniques such as visualization of the attention weights or the use of GNNExplainer^35^ to identify important subgraphs for classification can be performed. However, these may be difficult to interpret because they attempt to summarize complex interactions between high-dimensional data at the scale of thousands of patches per WSI. The complexity of such visualizations makes them difficult to understand and highlights the need for a simplified visualization.

Because similarity-based distances between patch-level GNN embeddings reflect higher-order connectivity and perceptually similar histological information, topological methods (such as Mapper) can compress this data to its essential structures while revealing the most salient aspects. For a given WSI, Mapper operates on the resulting point cloud of the patch-level GNN embeddings to first project the points to a lower dimensional space via techniques such as PCA, UMAP or NCVis ^36^ (referred to as *Projecting, f*). Once the data is projected, it is separated into overlapping sets (*Covering*, *U*), the number of which determines the resolution of the data summary. In each set, a *Clustering* algorithm (e.g. hierarchical clustering) is applied to the datapoints. The output of applying Mapper to this structure is a graph, where a node represents a cluster of WSI patches and an edge represent the degree of shared patches between the clusters^37^. This Mapper graph summarizes higher-order architectural relationships between patches and their shared histological information. In our framework, we refer to the nodes (collection of patches) as ROI, and the topological connectivity between the ROIs as their functional connectedness or “intermingling”. For instance, if a tumor ROI was connected to an ROI of the submucosa, we would say that the tumor has invaded (intermingled with) the submucosa. The degree of intermingling is quantified by the amount of overlap as defined by covering *U* and weighted by the incidence of cancer in each ROI. The expressiveness of this summary graph may be modified by selection of different *Filter*, *Cover*, and *Cluster* parameters which allows the user to interrogate ROIs in the WSI at different scales (a degree of flexibility beyond that of currently existing GNN pooling operations). We implemented Mapper using the *Deep Graph Mapper* implementation; however, python-based *Kepler-Mapper* and *giotto-tda* also present software solutions that may be readily employed ^37–39^.

#### 2.2.5. Associating ROI Connectivity with Clinical Outcomes

Once ROIs have been extracted using Mapper, measures of functional relatedness between the regions may be correlated with slide-level clinical outcomes. In our simple use case, we developed a Tumor Invasion Score (TIS) that measures the degree of overlap between the tumor and an adjacent tissue region. To construct this score, we first decompose each ROI into a vector encoding the frequency of each predicted tissue sub-compartment, 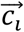 (counts of patch class assignment). The amount of overlap, as learned by Mapper’s *Cover* operation, between two ROI (*ROI*_*i*_ and *ROI*_*j*_, frequency vectors 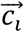 and 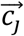 respectively) is *w*_*ij*_. The intermingling between different tumor sub-compartments for a pair of ROI may be expressed as 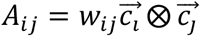. Given Mapper graph *G*, with edge-list *E*, (*e*_*ij*_ ∈ *E* the final pairwise associations between the regions are given by: 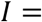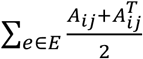. To measure tumor invasion/infiltration of the surrounding sub-compartments, we select the row of this matrix corresponding to the tumor: 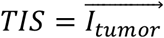 These vectors may be stacked across patients to form a design matrix, 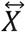, which may be associated with binary or continuous outcomes, 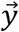. Here, we utilize Logistic Regression to associate *TIS* with cancer staging greater than 2 for the Colon samples, and positive Lymph Node status for Lymph Nodes.

### 2.3. Experimental Details

As proof of concept, we first pre-trained colon (comprised of epithelium, submucosa, muscularis propria, fat, serosa, inflammation, debris and cancer) and lymph node (fat, lymph node, cancer) classification networks with 10-fold cross validation (partitioning 10 separate training (82%), validation (8%) and test (10%) sets), evaluated using the area under the receiver operating curve (AUC/AUROC) and a weighted F1-Score. We extracted features from the penultimate layer of a ResNet50 neural network for about 2.7 million images per fold (26.9 million embedding extractions across 10-folds), using the pretrained network and separately from an ImageNet-pretrained model. After extraction of image features, we constructed graph datasets through calculation of the spatial adjacency (k-nearest neighbors) between the patches and storing node level embeddings into the attribute matrix. We created and trained a GNN that featured four graph attention layers, interspersed with ReLU activation functions ^40^ and DropEdge layers, followed by one layer of DropOut and finally a linear layer with SoftMax activation for node level prediction (**Figure 2A**). Models were generated using the pytorch-geometric ^28^ framework and trained using Nvidia v100 GPUs. For each cross-validation fold, we updated the parameters of our GNN through backpropagation of Cross-Entropy loss for node level classification on the training slides, while evaluating the potential to generalize on the validation set of *Whole Slide Graphs* (WSG) through evaluation of the F1-score. We saved the model parameters correspondent to the training epoch with the highest validation F1-score and extracted graph-node level embeddings and predictions on the validation and test sets of slides for each cross-validation fold. We refined patch-level predictions for all validation and test slides for the models with ImageNet-pretrained image features. To evaluate node-level tissue sub-compartment classification, we then calculated AUROC and F1-Score fit statistics across test slides. Finally, we applied Mapper to extract ROIs and *TIS* for all test slides (**Figure 2B**), of which 10-fold cross validation was applied over a non-penalized Logistic Regression model to estimate concordance with tumor stage and lymph node status. We also fit similar models to the frequencies of assignment of tissue sub-compartments for each case and combined relative frequencies of tissue sub-compartments with their *TIS* values to yield a final model. To evaluate the parsimony of the logistic regression model for alignment with our expectation that tumor infiltrating the fat corresponds to a high stage, we fit a generalized linear mixed effects model to the *TIS* scores, clustered by patient, and inspected the regression coefficients for quantitating the nature of these functional relationships.

**Fig. 2.**
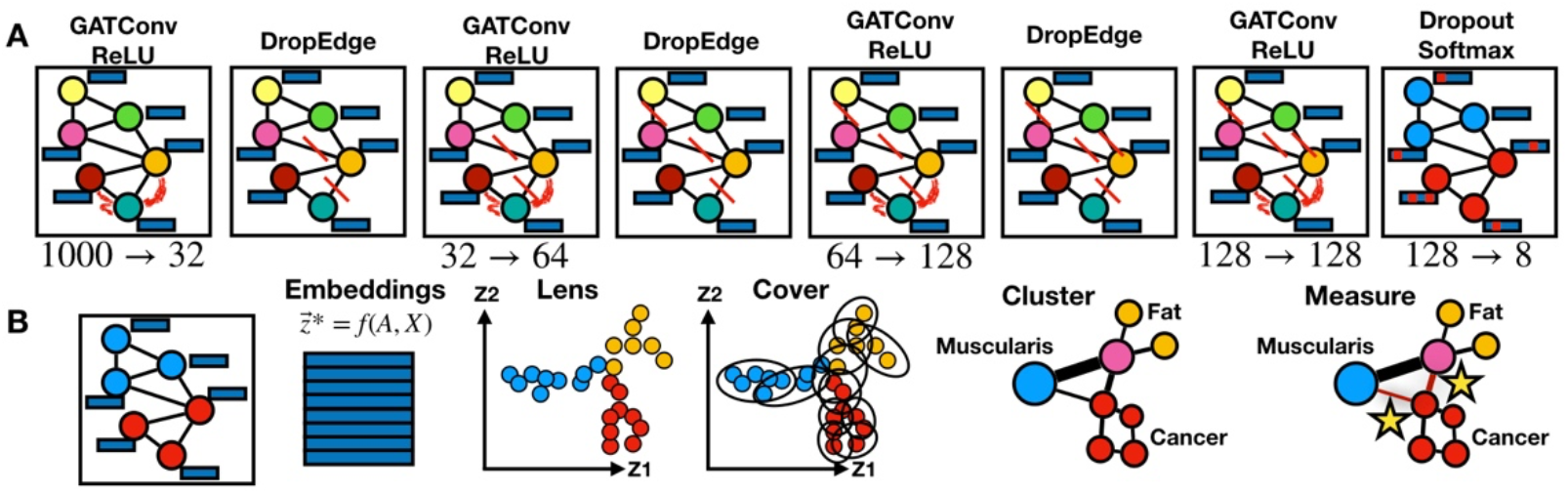
Methods: a) Neural network architecture for node classification experiment; 1000-d patch-level embeddings pass through graph attention convolutions, ReLU and DropEdge layers which alter dimensionality of patch embeddings while routing information from neighbors; attention between blue node and neighborhood is characterized using red curves; pruned edges are portrayed using red lines; b) once GNN classification model has been fit, GNN embeddings are extracted; lens function projects them to lower-dimensionality; patches are *covered* and *clustered* to reveal high-level measurable relationships between muscularis propria, fat and cancer

## 3. Results

### 3.1. Patch-Level Classification and Embeddings

An acceptable patch-level classification performance indicates room to further interrogate the slides for functional relationships between patches. In **Table 1**, we present 10-fold CV AUROC and F1-Score statistics on held-out test slides for patch level classification. The pretrained CNN for colon segmentation yielded moderately low performance metrics, while pretraining for lymph node yielded much higher scores. After feature extraction and training of the GNN taking into account the information from neighboring patches, scores increased substantially. Pretraining the CNN on Colon-specific targets had little impact on the classification model after fitting the GNN, suggesting that information contained from the patch surroundings is sufficient to contextualize that particular patch, or that the pretrained ImageNet is generalizable enough to histology images. Inspection of the patch-level embeddings (**Figure 3**), further corroborate that the original CNN does little to delineate the different classes of colon tissue, while the GNN embeddings demonstrate clear separation between these sub-compartments.

**Table 1.** GNN node classification results for colon (n=172 slides; 2,116,396 images) and LN (n=84; 570,326 images); averaged across slides; confidence assessed via 1000-sample non-parametric bootstrap

| 10-Fold CV (n=256) Node Classification | AUC ± SE | F1-Score ± SE |
| --- | --- | --- |
| Colon CNN-Only | 0.75±0.0054 | 0.43±0.0079 |
| Colon GNN ImageNet | 0.95±0.0026 | 0.81±0.006 |
| Colon Prediction Refinement | n/a | 0.83±0.0063 |
| Colon GNN Pretrained | 0.96±0.0031 | 0.82±0.0074 |
| LN CNN-Only | 0.91±0.0069 | 0.8±0.013 |
| LN GNN ImageNet | 0.96±0.0067 | 0.89±0.014 |
| LN GNN Pretrained | 0.97±0.0049 | 0.9±0.014 |

**Fig. 3.**
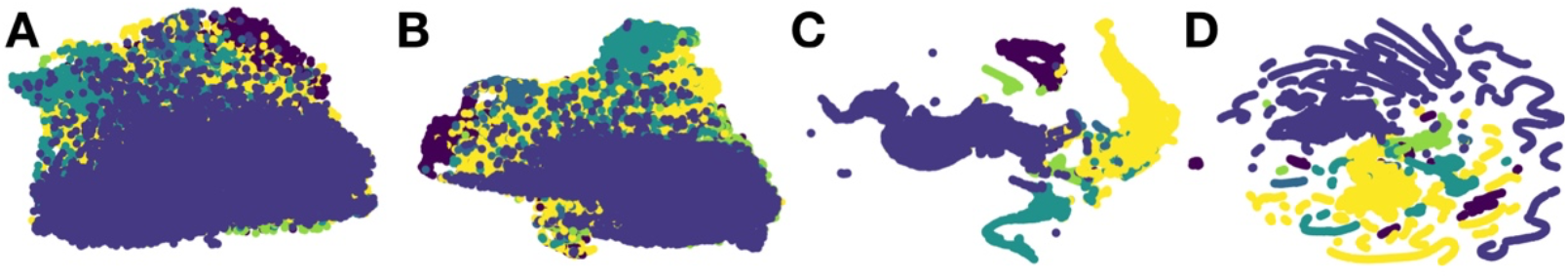
UMAP projection of penultimate layer of neural network for one select colon slide; nodes colored by true sub-compartment; a) ImageNet-pretrained CNN embeddings of patches; b) colon-pretrained CNN embeddings of patches; c) updated GNN patch embeddings after ImageNet extraction; d) updated GNN patch embeddings after colon-pretrained CNN extraction

### 3.2. Tumor Staging via Mapper Derived Invasion Scores

Figure 4 illustrates the extracted Mapper graph of representative low stage and higher stage slides. As compared to the lower stage slide, the *TIS* score derived for the higher stage indicates higher intermingling of tumor with regions of fat and is confirmed by pathologist annotations. We extracted an average of 32 ROIs from each WSI (range 5 – 155 ROI). Inspections of ROIs indicated some with clusters of tissue finely localized to histological tissue layer, and a few ROI that went undetected by the initial pathologist inspection (e.g. pockets of inflammation and tumor seeding in the fat).

**Figure. 4.**
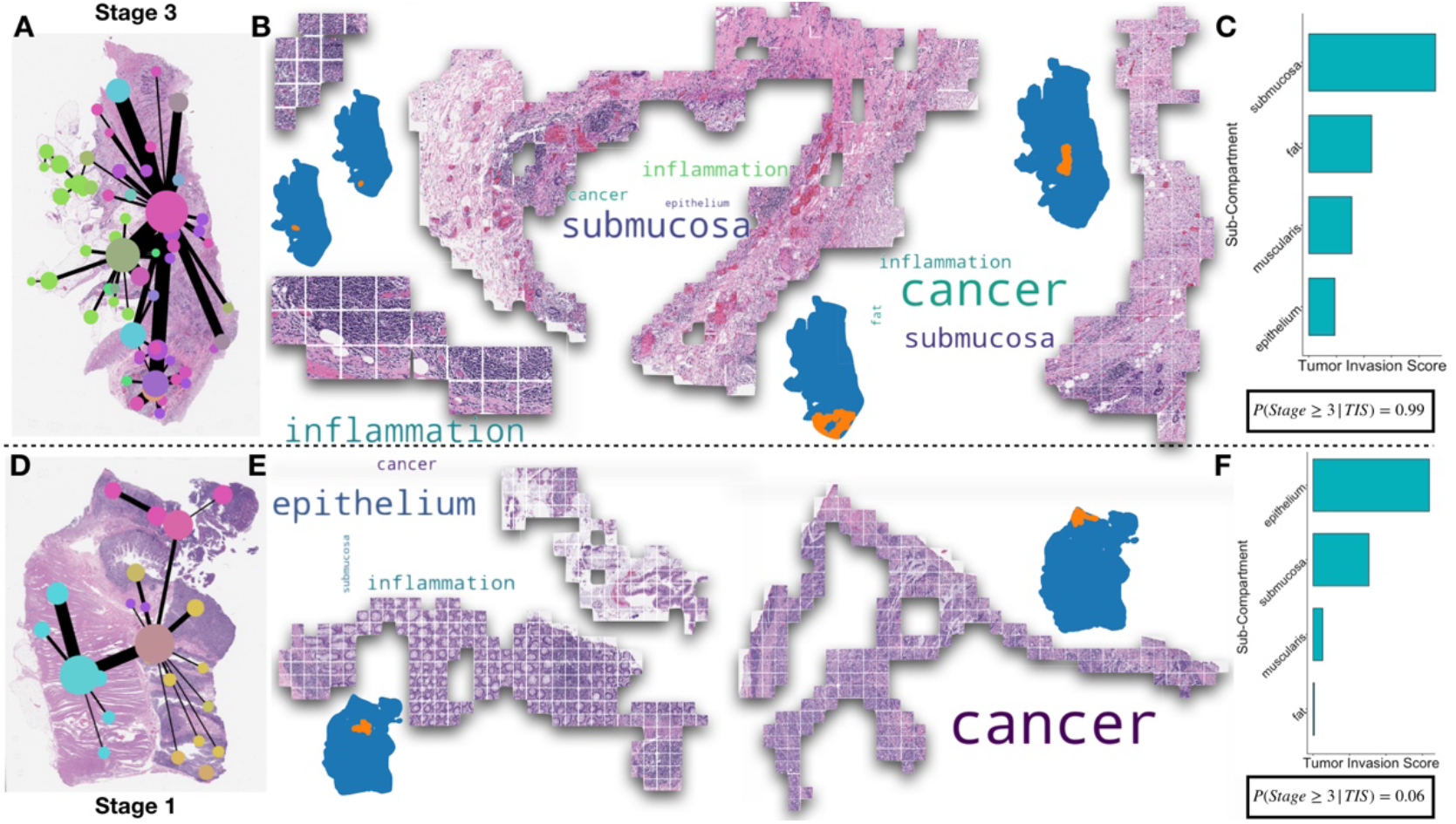
Example Topological Feature Extraction on two Colon slides; a) Mapper visualization of *WSG* of a stage 3 tumor; each vertex corresponds to an ROI, placement of the vertex reflects center of mass and thickness of edge connecting two points reflects topological overlap; b) example of four ROIs from the stage 3 slide; image patches are stitched back together; location is depicted in slide; composition of ROI is denoted with word clouds, where size of word is proportional to percentage makeup of ROI; c) actual *TIS* score for slide, prevalent invasion in the submucosa, fat and muscularis to reveal deep invasion; actual score from classifier gives 99% probability of advanced stage; d) Mapper visualization for stage 1 slide; e) left-most ROI demonstrates epithelial crypts with inflammation in lower right pocket; f) actual reported *TIS* score denotes invasion of epithelium with 6% probability of advanced stage

*TIS* scores correlated very well with Tumor staging. Ten-fold CV AUC was 0.91 for advanced Colon cancer staging and 0.92 for positive Lymph Nodes (**Table 2**). The frequency of sub-compartment instance and tumor invasion was also able to predict cancer stage when considered in isolation. Taken together, *TIS* and compartment localization achieved a higher AUC score, which speaks to the complementary information that each approach was able to provide to form a more complete picture of tumor progression. From the *TIS* scores, we were able to derive odds ratios (*OR*; measure of association between exposure and outcome, greater than one indicates adverse risk) as to their relation to tumor staging using linear mixed effects models (clustered on individual). As expected, fat interaction was highly associated with progression to a stage 3 or higher. Importantly, invasion of the muscularis propria, an adjacent and superficial region to the fat, had a statistically significant odds ratio commensurate with its depth in the colon.

**Table 2.** Ten-Fold AUROC statistics for unpenalized logistic regression prediction model on held out test data across all slides for colon (n=172) and LN (n=84); three head columns indicate whether advanced staging was predicted using aggregates of colon sub-compartment assignments (Region Counts); invasion (Tumor Invasion Scores); or Both to lend complementary information; models with main effects and interactions were considered; confidence assessed via 1000-sample non-parametric bootstrap

| AUC (10-Fold CV) | Region Counts Only |  | Tumor Invasion Scores Only |  | Both |  |
| --- | --- | --- | --- | --- | --- | --- |
| Stage > 2; LN Positive | Main Effects | Second Order Terms | Main Effects | Second Order Terms | Main Effects | Second Order Terms |
| Colon GNN Pretrained | 0.89±0.028 | 0.85±0.031 | 0.84±0.032 | 0.89±0.028 | 0.91±0.022 | 0.85±0.031 |
| Colon GNN Not Pretrained | 0.89±0.027 | 0.88±0.028 | 0.86±0.029 | 0.89±0.027 | 0.91±0.023 | 0.88±0.028 |
| LN GNN Pretrained | 0.76±0.077 | 0.88±0.046 | 0.88±0.046 | 0.76±0.077 | 0.88±0.046 | 0.88±0.046 |
| LN GNN Not Pretrained | 0.89±0.051 | 0.92±0.039 | 0.92±0.034 | 0.89±0.051 | 0.92±0.037 | 0.92±0.039 |

**Table 3.** Taking into account clustering on the patient level, odds-ratios derived from GLMM (ICC=0.21, n=172) fit on *TIS* scores derived from GNN that utilized colon-pretrained CNN embeddings; odds ratios indicate risk of advanced progression given tumor invasion of region

| *Stage* ≥ 3 |  |  |  |
| --- | --- | --- | --- |
| *Predictors* | *Odds Ratios* | *CI* | *p* |
| (Intercept) | 0.52 | 0.29 – 0.93 | **0.027** |
| *TIS: Epithelium* | 0.82 | 0.43 – 1.56 | 0.539 |
| *TIS: Fat* | 7.54 | 2.93 – 19.38 | **<0.001** |
| *TIS: Muscularis* | 1.68 | 1.02 – 2.77 | **0.043** |
| *TIS: Serosa* | 1.43 | 0.33 – 6.28 | 0.632 |
| *TIS: Submucosa* | 1.23 | 0.57 – 2.63 | 0.597 |

## 4. Discussion

Graph Neural Networks are increasingly promising approaches for studying WSI (and other gigapixel scale images) at multiple scales of inference through propagation of patch-wise information. However, when employing GNNs, the route of propagation often becomes obfuscated by the sheer quantity of patches being studied. This, in turn may make it difficult for researchers, clinicians or biologists to accept or understand these graph neural network technologies and their predictions. However, the compartmentalized and repetitive nature of tissue means that histology images can be greatly simplified via grouping of spatially adjacent subimages with perceptually similar and complementary input features. We have introduced methods from TDA to capture and reduce these motifs. In colon histology, we distilled information across WSI to better quantitate how the intermingling of different tissue sub-compartments inform disease stage. These results warrants investigation of other spatially driven processes, such as identifying ROIs correspondent to spatial transcriptomics^41^ and integration with high-dimensional omics data types.

Mapper has proven to be a useful TDA tool for elucidating high-level topology of the WSI. However, Mapper is highly dependent on the *Filter* function, *Cover* and *Cluster* parameters and algorithms to generate a topological map. While these features offer flexibility to study the WSI at multiple resolutions, which includes expanding the large range of ROI extracted, full exploration of the parameter space to identify an ideal range of parameters for Mapper graphs for the slide in study are beyond the scope of this work.

We also assessed the impact of domain-specific pre-training of a CNN on the resulting GNN predictions. Our preliminary results showed negligible impact on GNN accuracy. Integration of signal from the surrounding tissue context via GNNs may therefore be sufficient to overcome domain differences between histology images and real-world images (ImageNet). Further experimentation is needed on more nuanced examples to test this hypothesis.

There are a few limitations to our study. We assumed that GNNs are able to adequately capture patch-level information and their surrounding tissue architecture. The accuracy of our model was constrained by relatively coarse physician annotations that tended to ignore small structures like veins in the fat region of the lymph nodes, or small pockets of inflammation bounded by other tissue compartments, thereby reducing the accuracy of the model. However, inspection of regions with high uncertainty and label propagation allowed for correction of some of these issues. We also acknowledge the possibility of bias in given cross-validation folds. While we stratified the slides by whether they were representative of high or low stage, slides may contain different macroarchitectural features, and may, for instance, be completely devoid of serosa (which is only present in certain regions of the abdomen), which made it difficult to predict its presence. The colon WSI sections were analyzed from 36 patients (representing 256 slides). We acknowledge that there were repeated measurements taken across different slides from the same patient, the results of these sections may be correlated. While in our final inference on the *TIS* scores we account for this using mixed effects modeling, extracting samples from different patients would have been preferred to reduce cluster-level effects (data and pathologist time allowing). Due to the number of free parameters, we did not perform robust hyperparameter scans over the GNN.

In the future, we intend to utilize extracted GNN features contained within our ROIs to better identify the core topological structures that form a pathologist’s understanding of a slide^18^. Simplicial complexes represent series of points, lines, triangles and higher-dimensional tetrahedra. Persistence diagrams discover topological features in the form of simplicial complexes that persist over wide changes in proximity between points. These approaches can be readily applied to GNN embeddings to establish “barcodes” of various ROIs contained within the slide^42,43^, which may be used to supplement existing efforts to hash WSI to further assess the composition of other slides by the presence of characteristic topologies^44^. In addition to utilizing persistence based TDA methods, we aim to apply the aforementioned methods to GNN embeddings after applying graph pooling layers to identify topology and ROIs which may be related to molecular targets of interest, dense omics profiles and unlabeled clusters of tissue.

## 5. Conclusion

As multimodal deep learning approaches become increasingly important, GNNs are emerging as an attractive modeling tool for WSI representation where proper integration and association with slide-level outcomes is required. Conveniently, these approaches learn to identify key information pathways which may be simplified and visualized using TDA tools such as Mapper. Our method, *WSI-GTFE*, presents a framework from which to flexibly summarize the key insights acquired from fitting any GNN model to histological data. We hope that topological methods continue to see usage and integration with their deep learning graph counterparts for WSI level histological analyses given the benefits they provide in terms of model interpretability, quantitation of tissue compartment interaction, and potential for new biological discovery and disease prognostication.

## 6. Acknowledgements

*Preprint of an article published in Pacific Symposium on Biocomputing © 2021 World Scientific Publishing Co., Singapore, http://psb.stanford.edu/*

## Footnotes

* Software available on GitHub at the following URL: https://github.com/jlevy44/WSI-GTFE

